# BPP43_05035 is a *Brachyspira pilosicoli* cell surface adhesin that weakens the integrity of the epithelial barrier during infection

**DOI:** 10.1101/2024.04.06.584567

**Authors:** Anandi Rajan, Pablo Gallego, Brendan Dolan, Piyush Patel, Chinmay Dwibedi, Ana S. Luis, Sergio Trillo-Muyo, Liisa Arike, Sjoerd van der Post, Magnus Simrén, Thaher Pelaseyed

## Abstract

The anaerobic spirochete *Brachyspira* causes intestinal spirochetosis, characterized by the intimate attachment of bacterial cells to the colonic mucosa, potentially leading to symptoms such as diarrhea, abdominal pain, and weight loss. Despite the clinical significance of *Brachyspira* infections, the mechanism behind the interaction between *Brachyspira* and the colonic epithelium is not known. In this study, we characterized the molecular mechanism of *B. pilosicoli*-epithelium interaction and its impact on the epithelial barrier during infection. Through a proteomics approach, we identified BPP43_05035 as a candidate *B. pilosicoli* adhesion protein that mediates bacterial attachment to cultured human colonic epithelial cells. The crystal structure of BPP43_05035 revealed a globular lipoprotein with a six-bladed beta-propeller domain. Blocking the native BPP43_05035 on *B. pilosicoli*, either with a specific antibody or via competitive inhibition, abrogated its binding to epithelial cells. Furthermore, the binding of BPP43_05035 to epithelial cells required surface-exposed host *N*-glycans. Proximity labeling and interaction assays revealed that BPP43_05035 bound to tight junctions, thereby increasing the permeability of the epithelial monolayer. Extending our investigation to human patients, we identified a downregulation of tight junction and brush border genes in *B. pilosicoli*-infected patients carrying detectible levels of epithelium-bound BPP43_05035. Collectively, our findings identify BPP43_05035 as a *B. pilosicoli* adhesin that weakens the colonic epithelial barrier during infection.

## Introduction

The *Brachyspira* species, *B. aalborgi* and *B. pilosicoli*, are fastidious anaerobic bacteria that cause human colonic spirochetosis (*1*). A higher prevalence of *Brachyspira* species has also been reported in patients diagnosed with irritable bowel syndrome (IBS) with diarrhea (*2, 3*). Many enteropathogenic bacteria specialize in intimate binding to the intestinal epithelial cells of their host. Generally, bacteria gain access to the epithelial surface by breaking down host defense systems such as mucus and glycocalyx (*4*), followed by the deployment of cell surface components such as adhesins that latch onto host cells. In known instances, pathogens inject effector proteins that module host cell function to accommodate bacterial attachment, invasion, replication, and dissemination (*5, 6*).

*B. aalborgi* manifests an intimate interaction with colonic epithelial cells by establishing a perpendicular end-on attachment to the intermicrovillar space of the apical brush border (*7*). By contrast, *B. pilosicoli* is primarily associated with the inner mucus layer of the colon, where it can influence the underlying colonic epithelium (*3*). However, the molecular mechanism underlying the interaction of *B. pilosicoli* with the colon mucosa, especially when the mucus barrier is compromised, remains elusive.

We hypothesized that *B. pilosicoli* uses an undefined outer membrane protein (OMP) to engage specific host proteins on the apical surface of colonic epithelial cells. To test our hypothesis, we designed a proteomics-based discovery approach to identify OMPs involved in *B. pilosicoli* attachment to cells. We identified BPP43_05035, a globular six-bladed beta-propeller lipoprotein, which bound to cultured colonic epithelial cells through surface-exposed *N*-glycans. Notably, BPP43_05035 bound to cellular tight junctions where it triggered increased junction permeability. Finally, proteomic analysis of human colonic biopsies detected BPP43_05035 in patients infected with *B. pilosicoli* and further correlated its occurrence to transcriptional alterations in genes regulating tight junctions and the brush border in infected patients.

## Results

### Identification of BPP43_05035 as a putative *B. pilosicoli* adhesin

The human colorectal adenocarcinoma Caco-2 cell line was used as a model for exploring *B. pilosicoli* adhesins that participate in the interaction with the colonic epithelium. Caco-2 cells differentiate into a polarized epithelial monolayer defined by tight junctions along the course of 14-21 days (*8*). Initial observations under microaerophilic conditions confirmed the adhesion of cultured *B. pilosicoli* P43/6/78 to differentiated Caco-2 cell monolayers, preferably to cellular junctions (**Figure 1A**). Serial optical sections showed a perpendicular interaction between *B. pilosicoli* and the surface of Caco-2 cells (**Figure 1B**). Subsequently, we aimed to identify candidate OMPs that participate in this bacteria-host interaction. Viable *B. pilosicoli* were biotinylated with a negatively charged membrane-impermeable biotin reagent to specifically label OMPs exposed on the bacterial cell surface (**Figure 1C**). To specifically isolate adhesins, differentiated Caco-2 monolayers were incubated with a lysate of surface-biotinylated *B. pilosicoli* cells. Biotinylated OMPs were frequently observed on the apical cell surface and junctions between neighboring epithelial cells (**Figure 1D**). After successive washes, cell-bound OMPs were isolated by streptavidin affinity purification and analyzed by mass spectrometry. Proteins were identified by searching against the *B. pilosicoli* P43/6/78 protein database (Proteome ID: UP000010793), containing 2,208 protein entries. Search results were filtered for proteins present in all three replicates (Caco-2 + *B. pilosicoli* OMPs) but absent in the control samples (only Caco-2 cells), yielding a total of 85 unique *B. pilosicoli* proteins (**Table S1**). To identify putative OMPs, we applied additional filtering by selecting proteins containing a bacterial signal sequence. Among the most abundant signal sequence-containing proteins, we identified BPP43_05035 (UniProt ID A0A3B6VYW6), Peptidoglycan-associated lipoprotein (Pal) (UniProt ID K0JMH4), and putative treponemal membrane protein (tmpB) (UniProt ID K0JN47) (**Figure 1E**). In UniProt (*9*), BPP43_05035 was annotated as a sialidase-like protein and Pal as containing an OmpA-like domain (*10*). Of the three candidates, BPP43_05035 was highly enriched on the surface of Caco-2 cells since it constituted 7% of the bound OMPs but less than 1% of the *B. pilosicoli* proteome (**Figure 1F**). To corroborate the properties of three proteins as adhesins, we expressed and purified each candidate in *E. coli*. A polyclonal antibody generated against intact *Brachyspira* (*11*) recognized all three recombinant proteins, indicating that they are exposed on the outer membrane of *B. pilosicoli* cells (**Figure 1G**). Next, recombinant BPP43_05035, tmpB, and Pal were bound to differentiated Caco-2 cell monolayers. In analogy with viable *B. pilosicoli* and the crude mixture of biotinylated OMPs, BPP43_05035 localized to cellular junctions and the brush border, while Pal and tmpB variably formed protein aggregates on the monolayers (**Figure 1H**). The attachment of BPP43_05035 to the Caco-2 cell surface was cell-specific since it was not observed in the human colorectal carcinoma HCT116 cells, which lack the ability to form tight junctions (*12*) (**Figure 1H**). Consequently, we focused our investigation on the functional role of BPP43_05035 in the adhesion of *B. pilosicoli* to human epithelial cells.

**Figure 1.**
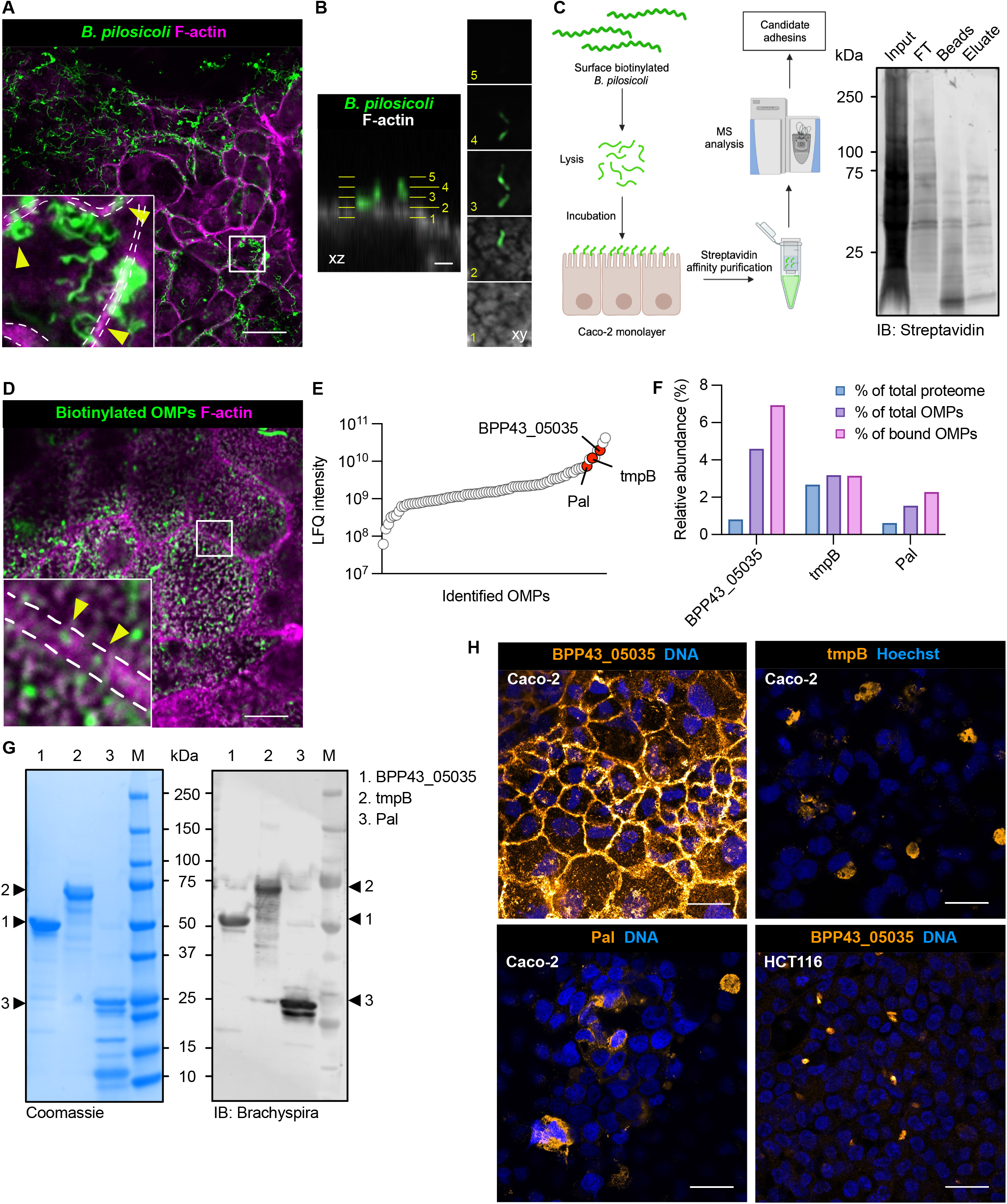
Isolation of *B. pilosicoli* outer membrane proteins that interact with Caco-2 cells. **(A)** Immunocytochemistry showing viable *B. pilosicoli* (green) attached to differentiated Caco-2 cells stained with Phalloidin (F-actin, magenta). The inset shows five times magnification. Dashed white lines mark the junctions between neighboring cells. Yellow arrowheads point to bacteria bound to cell junctions. Scale bar 20 µm. **(B)** Optical slices of *B. pilosicoli* bound to the apical surface of a Caco-2 cell. Scale bar 2 µm. **(C)** Experimental strategy for identifying *B. pilosicoli* outer membrane proteins (OMPs) that bind to Caco-2 cells (left panel). Immunoblot showing biotinylated proteins in the fractions input, flowthrough (FT), beads, and eluate, probed with Streptavidin. **(D)** Immunocytochemistry showing biotinylated OMPs (green) attached to differentiated Caco-2 cell stained with Phalloidin (F-actin, magenta). The inset shows five times magnification. Dashed white lines mark the junctions between neighboring cells. Yellow arrowheads point to biotinylated OMPs bound to cell junctions. Scale bar 10 µm. **(E)** Graph showing the label-free quantification (LFQ) of biotinylated *B. pilosicoli* OMPs bound to Caco-2 cells. Each point represents the LFQ intensity of one *B. pilosicoli* protein. Proteins are ordered by increasing LFQ intensity. BPP43_05035, Pal, and tmpB are highlighted in red. **(F)** Relative abundance of BPP43_05035, Pal, and tmpB in the entire *B. pilosicoli* proteome, in the sample with all the biotinylated OMPs, and in biotinylated OMPs bound to Caco-2 cells. **(G)** Coomassie gel and western blot with anti-*Brachyspira* antibody showing the separation of recombinant BPP43_05035, Pal, and tmpB using SDS-PAGE. Arrowheads point to each protein. M, molecular weight. **(H)** Immunocytochemistry showing the binding of BPP43_05035, Pal, and tmpB to Caco-2 cells, and binding of BPP43_05035 to HCT116 cells. Nuclear DNA is stained with Hoechst. Scale bar 20 µm.

### BPP43_05035 is a globular lipoprotein unique to *B. pilosicoli*

Having successfully purified recombinant BPP43_05035, we solved its crystal structure at 2.2 Å resolution (PDB ID: 7ZAO) (**Table S2**). BPP43_05035 consists of 463 amino acids, starting with an N-terminal signal sequence followed by a lipid attachment site (**Figure 2A**). The protein adopts a globular conformation comprising a six-bladed beta-propeller domain. (**Figure 2B**). BLAST similarity search excluding sequences belonging to *B. pilosicoli* showed a 25% sequence identity with a *Trypanosoma cruzi* trans-sialidase (**figure S1A, Table S2**). Further inspection using Distance Matrix Alignment (DALI) (*13*) identified the 50 closest structural homologs as various bacterial sialidases (**Figure 2C, figure S1B**). Structural alignment with the top homologs, NanA (2YA7) and NanI (2BF6), revealed an average root mean square deviation (RMSD) of 3.44 Å and 15% sequence identity (**Figure 2D**-**2E, figure S1C**). Additional alignment using Foldseek (*14*) identified six homologs classified as *B. pilosicoli* sialidases with an average RMSD of 1.17 ±0.21 Å and 35-37% sequence identity (**Figure 2C-D**). The closest structural homolog, the putative exo-alpha-sialidase EPJ72_11325, shared 70% sequence identity with BPP43_05035 (**Figure 2E-F**). In contrast, other putative *Brachyspira* sialidases such as EPJ79_11040 (*B. aalborgi*), BHYOB78_13105 (*B. hyodysenteriae*), and BPP43_01500 (*B. pilosicoli*) displayed high RMSD (4.67 Å) and low sequence identity (<20%) (**Figure 2C-E**). These findings suggest that BPP43_05035 is a unique *B. pilosicoli* protein with distant structural relatives among bacterial sialidases.

**Figure 2.**
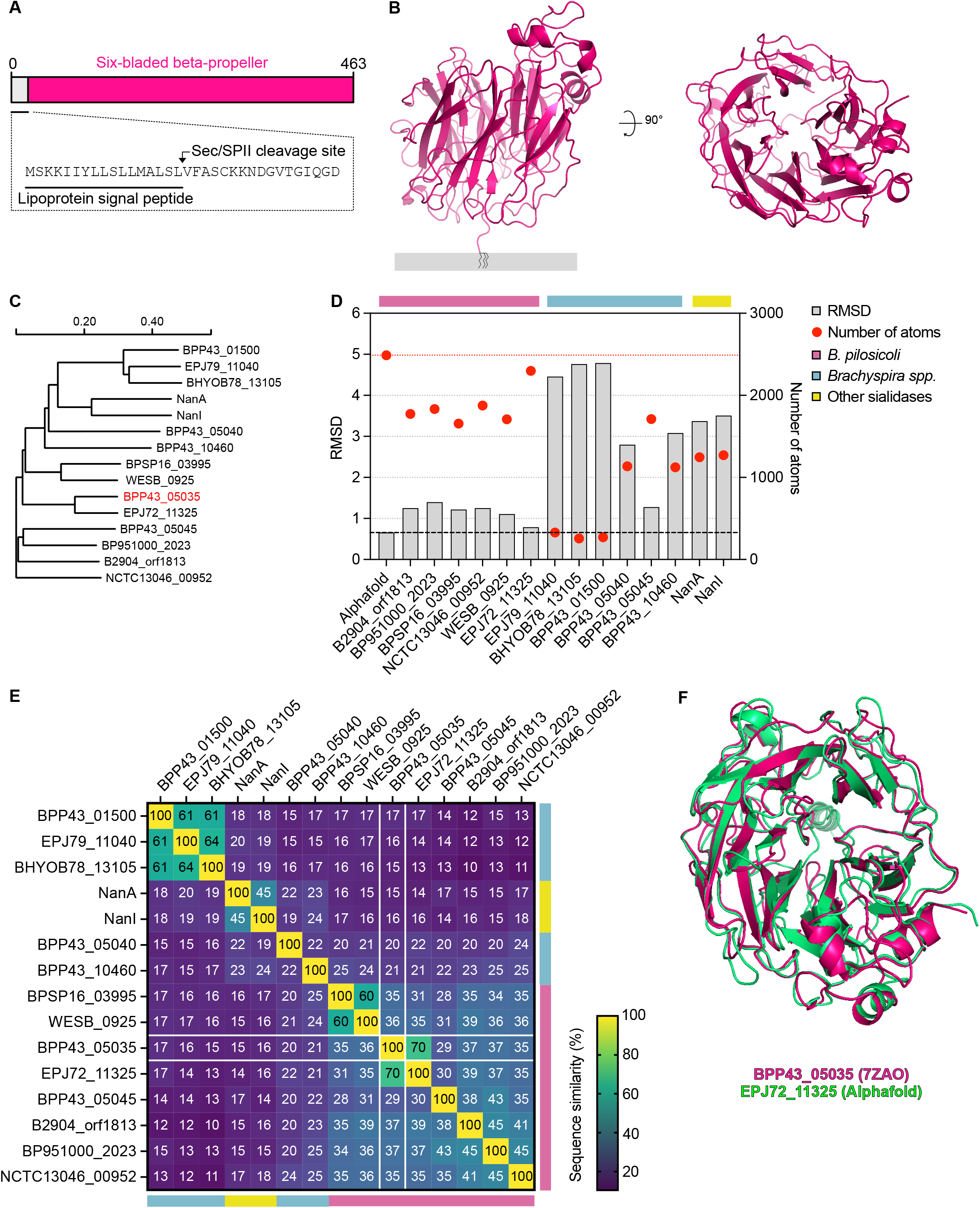
Structural characterization of BPP43_05035. **(A)** A schematic illustration of the full-length BPP43_05035. The signal sequence is underlined. The arrow points to the predicted position of the Sec/SPII cleavage site. **(B)** A cartoon representation of the crystal structure of BPP43_05035 (PDB ID 7ZAO) bound to a putative lipid moiety embedded in the outer membrane of *B. pilosicoli*. The right structure is a 90° projection of the left structure. **(C)** Phylogenetic tree showing the evolutionary relationship of the closest structural homologs of BPP43_05035. **(D)** Quantification of RMSD (left y-axis) and number of atoms included in each RMSD calculation (right y-axis) for the closest structural homologs of BPP43_05035 found in *B. pilosicoli*, *Brachyspira* spp., and other bacteria. RMSD is shown as gray bars. The number of atoms per RMSD calculation is shown as red circles. RMSD and number of atoms for the structural comparison between the empirical structure of BPP43_05035 and its Alphafold prediction are indicated by red and black dashed lines. **(E)** A percent identity matrix showing sequence identity between BPP43_05035 and structural homologs shown in (D). The Alphafold prediction is excluded from the matrix. The color scheme shows sequence similarity in percent (0-100%). **(F)** A cartoon representation showing an overlay of the structures for BPP43_05035 (magenta) and EPJ72_11325 (green).

The crystal structure of BPP43_05035 revealed a putative catalytic triad, consisting of a basic histidine (H221), an acidic glutamate (E282), and a nucleophilic serine (S340), positioned centrally in the protein (**figure S2A**). The occurrence of a catalytic triad is a hallmark of protein hydrolases (*15*). Thus, we assessed the activity of BPP43_05035 in hydrolysis assays with two classes of substrates. First, we used type II and III porcine gastric mucins (PGMs), containing 0.5-1.5% and <1.2% bound sialic acids, respectively, to evaluate if BPP43_05035 could hydrolyze sialic acids. A concentration of 10 μM BPP43_05035 was incubated with 0.5% of each PGM substrate across a pH spectrum of 5.5-8.5. Subsequent thin-layer chromatography revealed no detectable release of sialic acid from the PGM substrates (**figure S2B**), thereby negating the ability of BPP43_05035 to catalyze sialic acid hydrolysis. This finding was supported by the putative catalytic site of BPP43_05035 lacking structural resemblance to that of the neuraminidase domain in *Vibrio cholerae* NanH protein (**figure S2C**) (*16*). Next, we turned our attention to 4-nitrophenyl acetate and its derivatives, which serve as a class of standard substrates for measuring esterase activity. BPP43_05035 did not hydrolyze 4-nitrophenyl acetate at pH 5.5-8.5 (**figure S2D**) or longer butyl and octyl esters at pH 7 (**figure S2E**). Taken together, our observations indicate that BPP43_05035 does not possess any enzymatic activity against sialic acid or ester bonds.

### BPP43_05035 is required for the attachment of *B. pilosicoli* to Caco-2 cells

To investigate the localization of BPP43_05035 in *B. pilosicoli* cells, we generated a polyclonal antibody against the globular six-bladed beta-propeller domain. The antibody detected BPP43_05035 as distinct puncta on *B. pilosicoli*, including on the opposite ends of the spiral-shaped cell (**Figure 3A**). Next, we devised two independent strategies to assess if BPP43_05035 is required for the adhesion of *B. pilosicoli* to Caco-2 monolayers (**Figure 3B**). First, viable *B. pilosicoli* were pre-incubated with either the anti-BPP43_05035 antibody or a pre-serum control prior to binding the bacteria to the epithelial monolayers. Pre-incubation with the anti-BPP43_05035 antibody significantly inhibited the attachment of viable *B. pilosicoli* to the cell (**Figure 3C**). In the second strategy, we took advantage of the purified BPP43_05035 to block and saturate its binding sites on differentiated Caco-2 cells before incubation with viable *B. pilosicoli*. In analogy with the BPP43_05035-specific antibody, saturating BPP43_05035 binding sites with the recombinant protein significantly reduced the binding of *B. pilosicoli* to the epithelial cells (**Figures 3D**). Thus, obstructing endogenous BPP43_05035 on the bacterial cell or competitively blocking its corresponding binding site on target cells inhibited the adhesion of *B. pilosicoli* to human epithelial cells.

**Figure 3.**
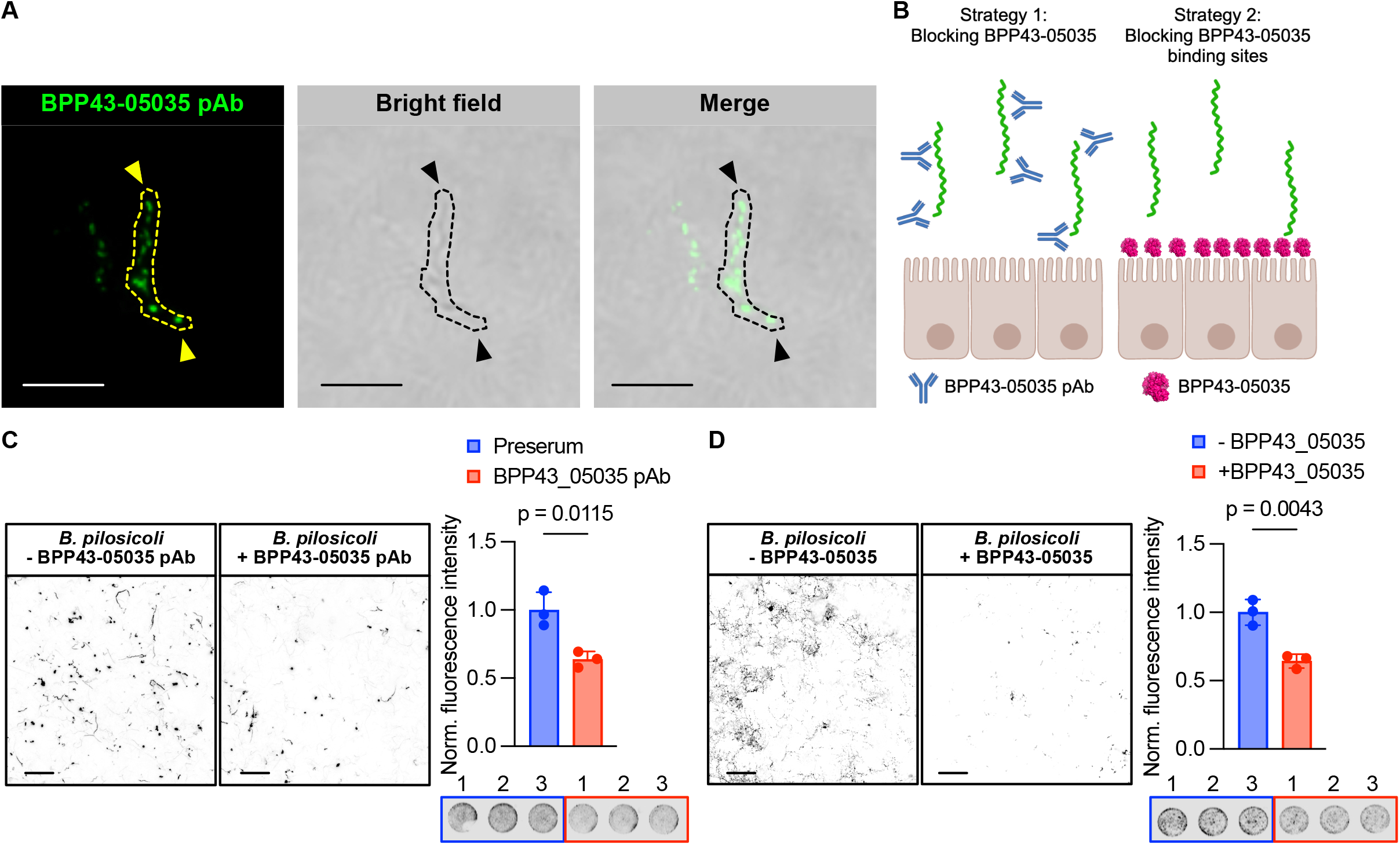
BPP43_05035 is required for the attachment of *B. pilosicoli* to Caco-2 cells. **(A)** Immunostaining of unpermeabilized *B. pilosicoli* with a polyclonal antibody raised against BPP43_05035. The confocal image is superimposed on a brightfield image. An individual bacteria cell is enclosed with a dashed yellow or black line. Arrows point to BPP43_05035 at the opposing ends of the bacterial cell. Scale bars 10 µm. **(B)** A schematic presentation of the two strategies, involving the BPP43_05035-specific antibody or recombinant BPP43_05035 protein, to inhibit the binding of *B. pilosicoli* to Caco-2 cells. **(C)** Immunocytochemistry of *B. pilosicoli* bound to Caco-2 cells after preincubation with the BPP43_05035 polyclonal antibody (left). Quantification of normalized fluorescent intensity of fluorescently labeled *B. pilosicoli* bound to Caco-2 cells after preincubation with the BPP43_05035 polyclonal antibody (right). The panel below the bar graph shows representative images of the binding assay in 96-well format. Scale bars 20 µm. n=3 for each group. Data are means ± SD. Significance was determined by unpaired t-test. **(D)** Immunocytochemistry of *B. pilosicoli* bound to Caco-2 cells after preincubation of Caco-2 cells with the recombinant BPP43_05035 protein (left). Quantification of normalized fluorescent intensity of fluorescently labeled *B. pilosicoli* bound to Caco-2 cells after preincubation of Caco-2 cells with the recombinant BPP43_05035 (right). The panel below the bar graph shows representative images of the binding assay in 96-well format. Scale bars 20 µm. n=3 for each group. Data are means ± SD. Significance was determined by unpaired t-test.

### BPP43_05035 binds to host cell surface *N*-glycans

To investigate the interaction of BPP43_05035 with the epithelium in a more complex cellular context, we turned to biopsies collected from the sigmoid colon of healthy individuals. Binding assays on histological sections revealed that BPP43_05035 bound to *N*- and *O*-glycan-rich cellular compartments, particularly the apical brush border covered with a membrane mucin-based glycocalyx (*17*) and goblet cells storing the gel-forming mucin MUC2 (**Figure 4A**). To further evaluate the specificity of BPP43_05035 for colonic mucins, we assessed the binding of BPP43_05035 to colonic tissue sections of *Muc2^-/–^* mice. While BPP43_05035 bound to goblet cells in the mouse colon, the goblet cell-specific binding was completely abrogated in the *Muc2^-/–^* colon, which did not express Muc2 (**Figure 4B**). To distinguish between host *N*- and *O*-glycans conjugates as potential receptors for BPP43_05035, we screened the glycan specificity of BPP43_05035 using a comprehensive array of 562 distinct carbohydrate structures. While BPP43_05035 did not bind any *O*-glycans, we observed weak interactions with *N*-glycans at three different concentrations of BPP43_05035 (**Figure 4C-D, figure, S3, Table S3**). Subsequently, we investigated the role of host *N*-glycans in the binding of BPP43_05035 to epithelial cells using the *N*-glycosidase F, which cleaves between the innermost GlcNAc of N-linked glycans and asparagine in the glycoproteins. Compared to untreated cells, the binding of BPP43_05035 to differentiated Caco-2 cell monolayers was reduced after treatment with *N*-glycosidase F (**Figure 4E**), proposing that the interaction is primarily dependent on surface-exposed *N*-glycans.

**Figure 4.**
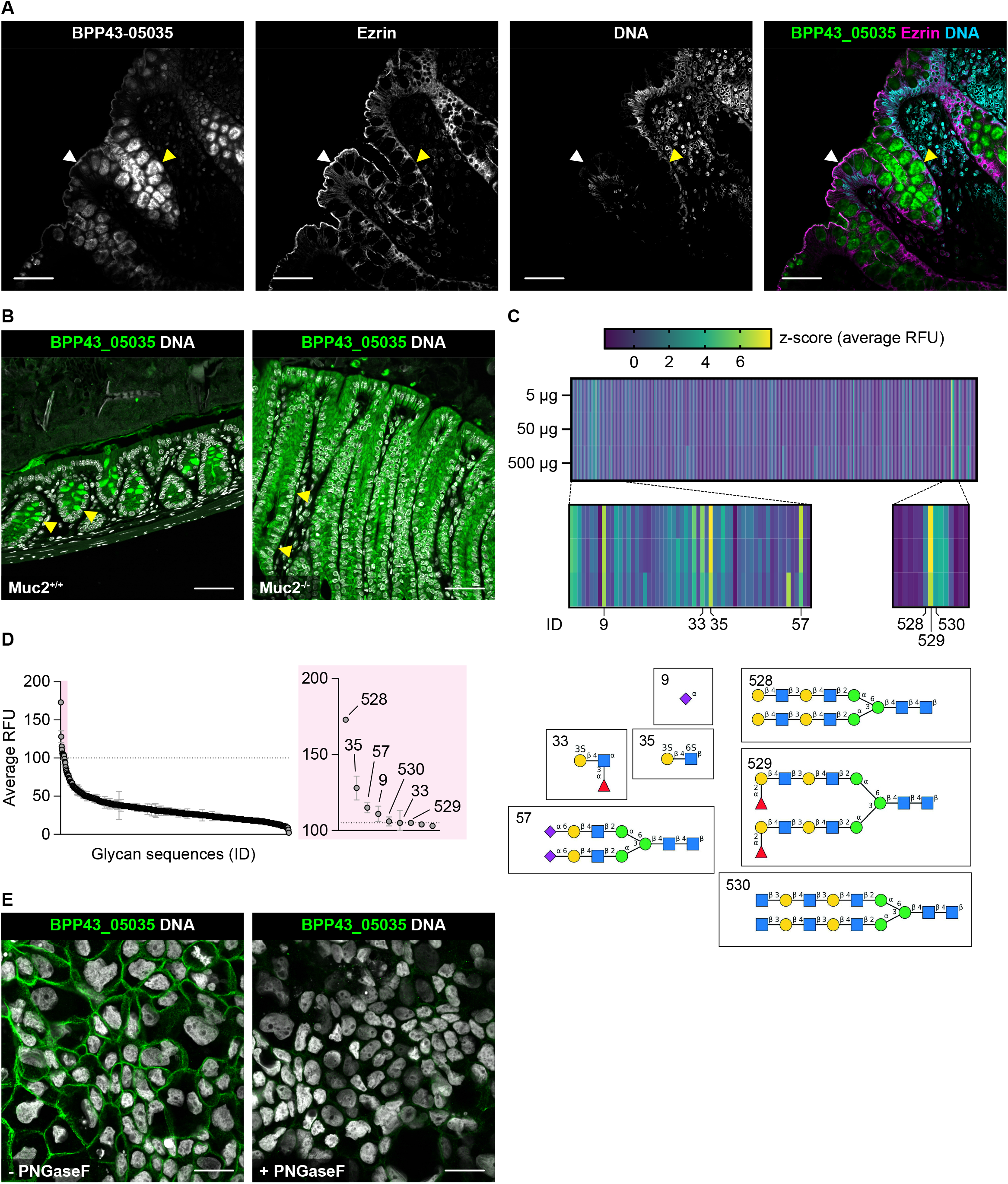
BPP43_05035 binds to human colonic biopsies and requires a surface-exposed host *N*-glycan. **(A)** Immunohistochemistry showing the binding of BPP43_05035 (green) to sections of human sigmoid colon stained for Ezrin (magenta) and nuclear DNA (blue). The white arrowhead points to the apical brush border. Yellow arrowhead points to goblet cell. Scale bars 50 µm. **(B)** Immunohistochemistry of binding of BPP43_05035 (green) to sections of the distal colon from *Muc2^+/+^* and *Muc2^-/^*^-^ mice. Nuclear DNA is shown in gray. Yellow arrowheads point to goblet cells. Scale bars 50 µm. **(C)** A heatmap showing the binding of BPP43_05035 (5-500 µg) to a glycan array of 562 glycans in 6 replicates (CFGv5.5). The glycan structures are ordered based on ID number 1-562. The color scheme represents z-scores of average relative fluorescence units (RFU). Structures for ID numbers with the highest z-score are shown below the heatmap. **(D)** A bar graph showing the average RFU for the binding of BPP43_05035 to the glycan array. The glycan structures represented in (C) are indicated with their corresponding ID numbers. **(E)** Immunocytochemistry showing the binding of BPP43_05035 (green) to untreated or *N*-glycosidase F-treated differentiated Caco-2 cells. Nuclear DNA is shown in gray. Scale bars 20 µm.

### BPP43_05035 interacts with proteins that regulate intercellular tight junctions

To identify the epithelial host receptor for *B. pilosicoli*, we used proximity biotin labeling by antibody recognition (*18*) to tag endogenous epithelial cell proteins using the bacterial cell as bait. Viable *B. pilosicoli* were bound to differentiated Caco-2 monolayers, followed by fixation, blocking, and probing for bacteria using the *Brachyspira*-specific antibody and a peroxidase-conjugated secondary antibody. All proteins in the proximity of bacterial cells were labeled with biotin tyramide in the presence of peroxide. Affinity purification with immobilized streptavidin captured the biotinylation of bacterial and epithelial proteins in the presence of primary and secondary antibodies and biotin tyramide (**Figure 5A**). A specific band, which was not detected with the *Brachyspira*-specific antibody, was observed at 37 kDa and shown by proteomic analysis to contain the actin-binding protein Shroom3, localized to apical junctional complexes (*19*), the junctional protein JUP, the actin-plasma membrane crosslinker EZR, and the Junctional adhesion molecule 1 (JAM-1, also called JAM-A or F11R), which localizes to tight junction complexes where it regulates the integrity of the epithelial cell barrier (*20*) (**Figure 5B, Table S4**). String analysis revealed that the labeled proteins participated in terminal web assembly (GO:1902896) and protein localization to bicellular tight junction (GO:1902396) (**Table S4**). In a parallel experiment, we sought to identify intracellular host proteins, which may interact with BPP43_05035. A co-immunoprecipitation assay using BPP43_05035 as bait in lysates of Caco-2 cells, followed by a mass spectrometric analysis of co-immunoprecipitated Caco-2 proteins separated by SDS-PAGE, identified proteins that interacted with BPP43_05035 but not with preserum-conjugated or unconjugated beads (**Figure 5C, Table S5**). The identified proteins included Actin (ACTB), Myosin-9 (MYH9), and Myosin regulatory light chain 12B (MYL12B, MLC2), which form a functional interaction network according to String DB (**Figure 5D**). KEGG pathway analysis of the BPP43_05035 interactome revealed pathways involved in tight junctions (hsa04530), regulation of the actin cytoskeleton (hsa04810), and bacterial invasion of epithelial cells (hsa05100) (**Figure 5E**). The BPP43_05035 interactome was annotated as Gene ontology cellular component brush border (GO:0005903), actin cytoskeleton (GO:0015629), and stress fiber (GO:0001725) (**Figure 5F**), which all are associated with tight junction function (*21*). In summary, two independent biochemical assays showed that BPP43_05035 associates with proteins that form and regulate cellular tight junctions.

**Figure 5.**
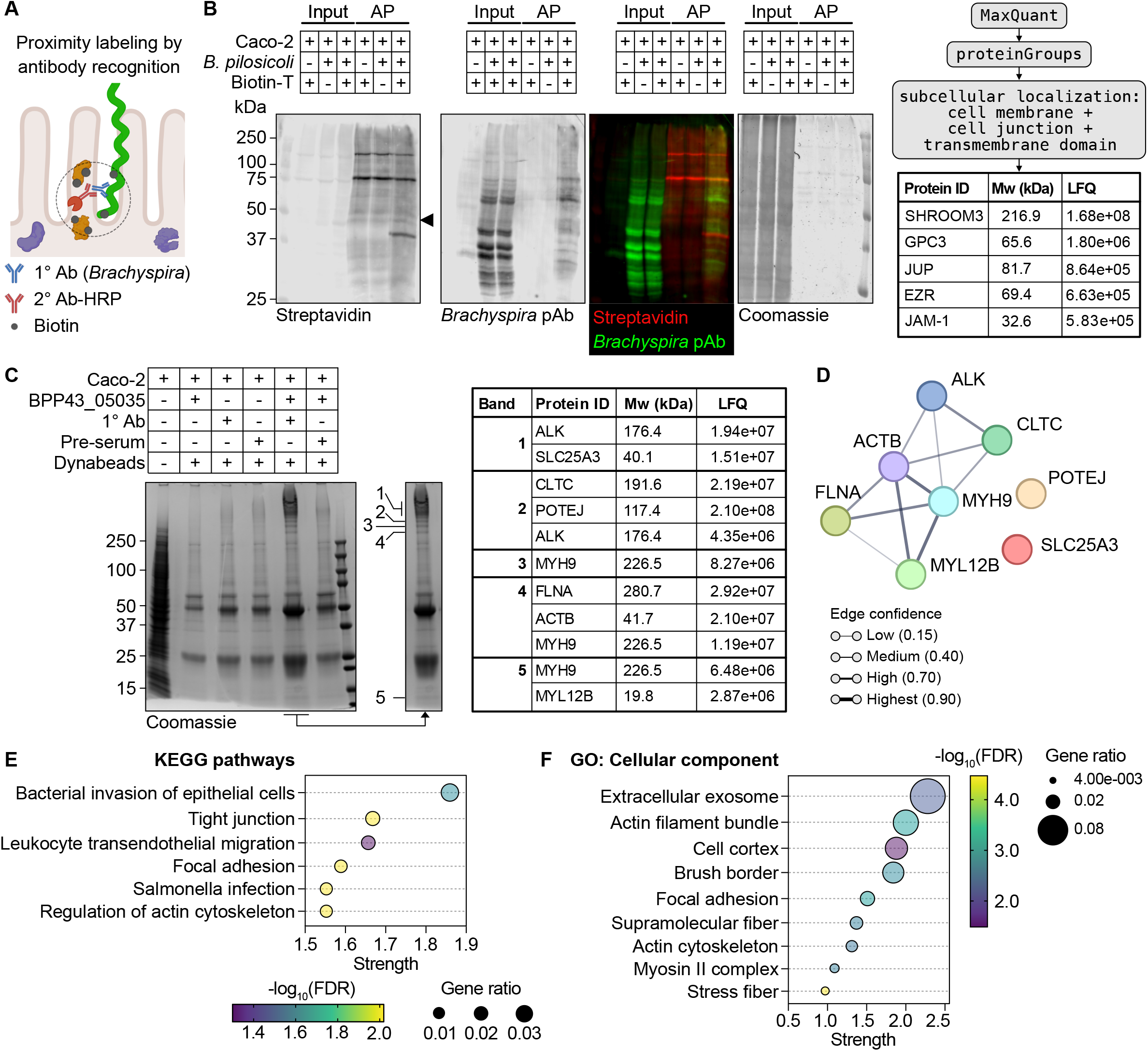
BPP43_05035 associates with proteins that regulate tight junctions. **(A)** A Schematic presentation of proximity labeling by antibody recognition using the *Brachyspira*-specific primary (1°) antibody and an HRP-conjugated secondary (2°) antibody. **(B)** A two-channel immunoblot representing the inputs and the biotinylated proteins isolated via streptavidin affinity purification (AP) after proximity labeling using B. pilosicoli as bait. Biotinylated proteins were probed by Streptavidin (red channel) and a *Brachyspira*-specific antibody (green channel). The Coomassie membrane stain shows the total protein amounts loaded in each lane. A description of the bioinformatic pipeline used to identify proximal proteins that localize to cell membranes and junctions or contain a transmembrane domain. The table shows the identified proteins, with their corresponding molecular weight (Mw) and label-free quantification (LFQ) intensity. **(C)** A Coomassie gel staining showing proteins co-immunoprecipitated with recombinant BPP43_05035 as bait. The lane of interest is also shown separately, and the protein bands analyzed by mass spectrometry are numbered 1-5. The table shows the identified proteins, with their corresponding molecular weight (Mw) and label-free quantification (LFQ) intensity, in each band. **(D)** STRING analysis of the identified proteins presented in C. **(E)** KEGG pathway analysis of the identified proteins in C. The color scheme shows the false discovery rate, calculated as -log_10_(FDR). The size of each point represents the gene ratio defined as the percentage of significant genes over the total number of genes in the pathway. The x-axis shows strength (enrichment effects), defined as the ratio between the number of identified proteins annotated within a term and the number of proteins expected to be annotated within the term in a random network of the same size. **(F)** Cellular component gene ontology analysis of the identified proteins in C.

### BPP43_05035 interacts with tight junctions and impairs epithelial barrier integrity

Tight junctions are multi-protein complexes that regulate the barrier integrity of the epithelium. Prompted by the association of BPP43_05035 with JAM-1 and modulators of tight junctions, we sought to understand the impact of BPP43_05035 on tight junction function. Treatment of Caco-2 monolayers with BPP43_05035 resulted in a profound change in the morphology of tight junctions stained with ZO-1, from typically convex curves to irregular curves that connect tricellular junctions (**Figure 6A**). Quantification of the staining profile of ZO-1 showed a significant loss of straightness between individual tricellular junctions (**Figure 6B**). To define the spatial distribution of BPP43_05035 in the monolayers, we analyzed its localization in relation to JAM-1, ZO-1, and an additional tight junction protein, Occludin (**Figure 6C-E**). Quantitative analysis of fluorescent intensities along transects across intercellular junctions revealed a notable correlation between the spatial distribution of BPP43_05035 and the selected tight junction proteins (**Figure 6C-E**). Based on the distinct overlap between BPP43_05035 and the tight junction proteins, we postulated that BPP43_05035 influences the permeability of tight junctions in live cells. Differentiated Caco-2 monolayers were incubated with either 100 μM of BPP43_05035 or viable *B. pilosicoli* cells, followed by measurement of transepithelial electrical resistance (TEER) to quantify the paracellular integrity maintained by tight junctions. The binding of BPP43_05035 to the monolayer resulted in a marked decrease in TEER compared to untreated cells (**Figure 6F**). Likewise, Caco-2 monolayers exposed to viable *B. pilosicoli* for an extended 30-hour period under microaerophilic conditions manifested a 60% reduction in TEER, compared to a mere 25% decrease in the untreated cells (**Figure 6G**) and the comparable attenuation of TEER from start to the end of the experiment was more pronounced in the *B. pilosicoli*-treated compared to the untreated cells (**Figure 6G**). Collectively, our observations provide evidence that BPP43_05035 binds to tight junctions and compromises the barrier integrity of epithelial monolayers.

**Figure 6.**
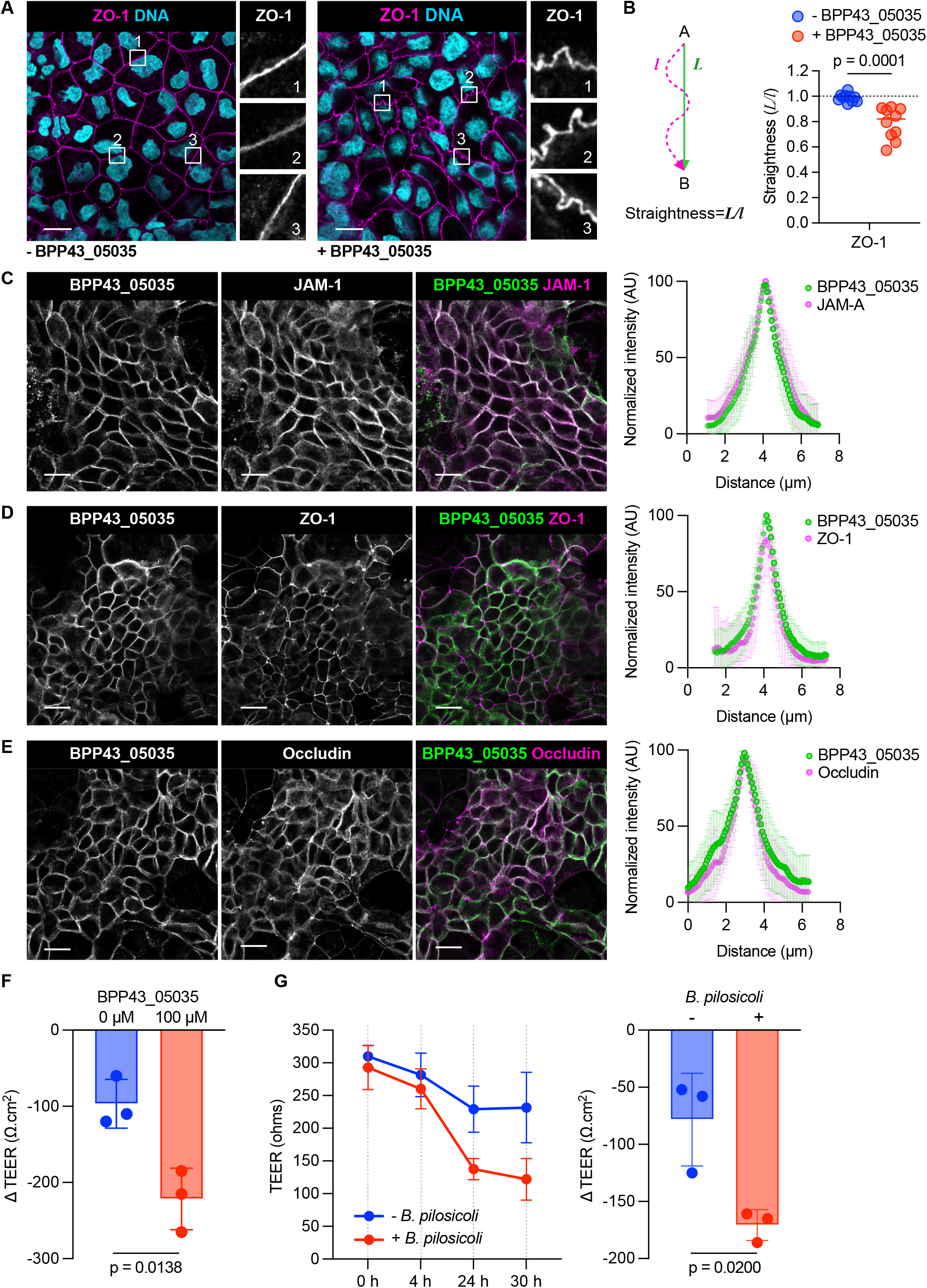
BPP42_05035 interacts with tight junctions and impairs tight junction barrier integrity. **(A)** Immunocytochemistry of differentiated Caco-2 cells, untreated or treated with BPP43_05035, and stained with ZO-1. Nuclear DNA is stained with Hoechst. Numbered insets represent 4x magnifications of the selected region. Scale bars 20 µm. **(B)** Quantification of tight junction morphology, defined as the staining profile of ZO-1 between two interconnecting tricellular junctions, in untreated and BPP43_05035-treated Caco-2 cells. n=10 measurements per group. Data are shown as mean ± SD. Significance was determined by unpaired t-test. **(C)** Immunocytochemistry of differentiated Caco-2 cells incubated with BPP43_05035 and stained with the tight junction protein JAM-1. The normalized intensity of BPP43_05035 in relation to JAM-1 along transects across intercellular junctions is shown to the left. n=15 measured transects per group. Data are shown as mean ± SD. Scale bars 20 µm. **(D)** Immunocytochemistry of differentiated Caco-2 cells incubated with BPP43_05035 and stained with the tight junction protein ZO-1. The normalized intensity of BPP43_05035 in relation to ZO-1 along transects across intercellular junctions is shown to the left. n=15 measured transects per group. Data are shown as mean ± SD. Scale bars 20 µm. **(E)** Immunocytochemistry of differentiated Caco-2 cells incubated with BPP43_05035 and stained with the tight junction protein Occludin. Normalized intensity of BPP43_05035 in relation to Occludin along transects across intercellular junctions is shown to the left. n=15 measured transects per group. Data are shown as mean ± SD. Scale bars 20 µm. **(F)** Quantification of the difference in TEER (ΔTEER), measured at the start and the end of the experiment, in untreated and BPP43_05035-treated Caco-2 cells. n=3 for each group. Data are shown as mean± SD. Significance determined by unpaired t-test. **(G)** Time course TEER measurement in Caco-2 cells, untreated or treated with viable *B. pilosicoli.* n=3 for each group and time point. Data are shown as mean ± SD. Quantification of the difference in TEER (ΔTEER), measured at the start and the end of the experiment. n=3 for each group. Data are shown as mean ± SD. Significance determined by unpaired t-test.

### Presence of BPP43_05035 in the human colon correlates with altered tight junction and brush border gene signatures

Previous studies detected *B. pilosicoli* in the sigmoid colon of a subset of patients diagnosed with IBS (*3*). Consequently, we performed transcriptomic analysis on biopsies collected from the sigmoid colon of uninfected individuals (healthy controls, n=5) and individuals with confirmed *B. pilosicoli* infection (participants with IBS, n=5), validated independently by quantitative real-time PCR (*3*) **(Table S6)**. Principal component analysis and unsupervised hierarchical cluster analysis differentiated the two patient groups based on genes belonging to tight junctions (GO:0005923) and the apical brush border (GO:0005903) (**Figure 7A**), demonstrating an overall downregulation of genes associated with the two cellular compartments (**Figure 7B, Table S7**). Notably, the majority (60%) of the genes contributing to PC1 were associated with tight junctions, whereas brush border and tight junctions genes contributed equally to PC2, suggesting that the differences between the two groups are mainly explained by alterations in the transcription of tight junction genes (**Figure 7C**). Next, we performed proteomic analysis on isolated intestinal epithelial cells collected from the sigmoid colon of each group. Expectedly, we identified a higher number of peptides belonging to *B. pilosicoli* in the infected group. The top 50 *Brachyspira* proteins included previously identified *B. pilosicoli* extracellular proteins (*22*) (**Figure 7D, Table S8**). Notably, BPP43_05035 was the most abundant *B. pilosicoli* protein in the epithelial proteome of the infected patients (**Figure 7E**), suggesting that BPP43_05035 is intimately attached to the colonic epithelium during infection. In summary, we provide the first experimental evidence revealing the critical role of the outer membrane protein BPP43_05035 in the adhesion of the spirochete *B. pilosicoli* to the human colonic epithelium. Our study suggests that BPP43_05035 is an integral component of *B. pilosicoli*’s attachment mechanism and a key determinant of *B. pilosicoli* colonization of the human colon.

**Figure 7.**
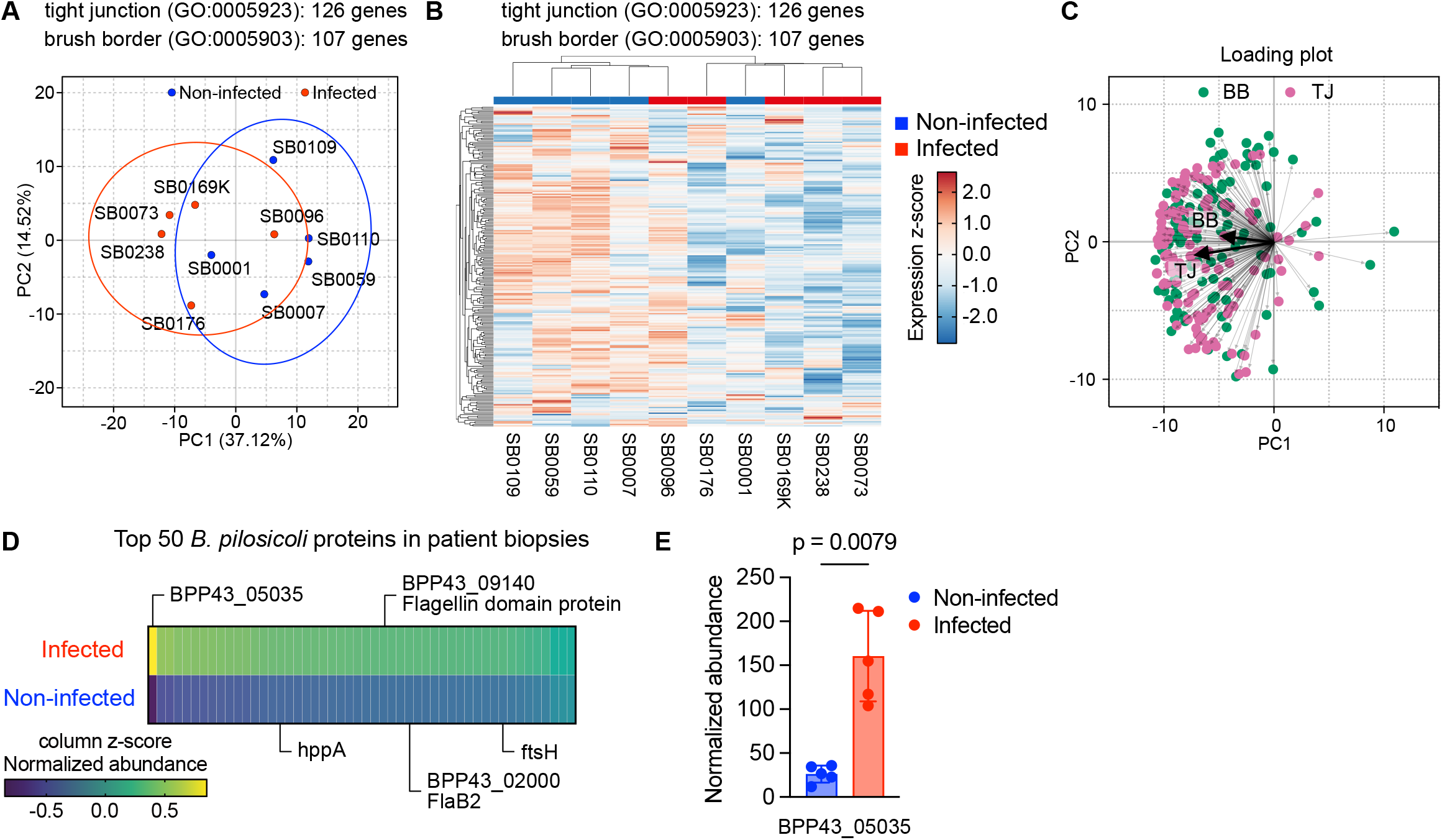
Detection of the epithelium of patients infected with *B. pilosicoli*. **(A)** Principal component analysis of sigmoid colon biopsies from non-infected and *B. pilosicoli*-infected patients, based on gene expression signatures belonging to GO:0005923 (tight junction) and GO:0005903 (brush border). **(B)** Heatmap showing expression z-score of gene signatures belonging to GO:0005923 (tight junction) and GO:0005903 (brush border) in the sigmoid colon of non-infected and *B. pilosicoli*-infected patients. **(C)** A loading plot showing the contribution of individual brush border (green) and tight junction (purple) genes to PC1 and PC2. The bold arrows show the median contribution of brush border and tight junction genes. **(D)** Top 50 *B. pilosicoli* proteins identified in biopsies collected from the sigmoid colon of non-infected and *B. pilosicoli*-infected patients. The color scheme shows the column z-score of normalized protein abundance. Specific proteins are labeled with names. **(E)** Quantification of the normalized abundance of BPP43_05035 in the sigmoid colon of non-infected and *B. pilosicoli*-infected patients. n=5 in each group. Data are shown as mean ± SD. Significance determined by unpaired t-test.

## Discussion

The most reliable clinical evidence of *Brachyspira* infection is the presence of spirochetes attached to the epithelial brush border in colorectal biopsy specimens. However, the molecular mechanisms by which *B. pilosicoli* adheres to the colonic mucosa are not understood. Previous analyses of the *B. pilosicoli* genome have failed to identify genes encoding virulence factors involved in attachment (*23*). In this study, we identified BPP43_05035 that mediated an *N*-glycan-dependent attachment of *B. pilosicoli* to colonic epithelial cells. BPP43_05035 was associated with cellular tight junctions and increased junctional permeability in *in vitro* cell cultures. Importantly, we detected BPP43_05035 in colonic specimens collected from *B. pilosicoli*-infected participants with impaired expression of tight junction and brush border genes.

BPP43_05035 is a *bona fide B. pilosicoli* OMP since it was isolated via surface biotinylation of viable bacterial cells grown under anaerobic conditions and detected by an antiserum against intact spirochetes. Moreover, an antibody raised against recombinant BPP43_05035 detected the endogenous protein at the extreme ends of unpermeabilized *B. pilosicoli* cells, corresponding to regions where the bacterium forms an invagination into the apical membrane of host cells. Since the genetic manipulation of *Brachyspira* species has not been made possible (*24*), the BPP43_05035-specific antibody was able to inhibit the binding of *B. pilosicoli* to epithelial cells, indicating that surface-exposed BPP43_05035 mediated the attachment of *B. pilosicoli* to host cells.

Extensive structural and sequence comparison with bacterial proteins revealed that BPP43_05035 is unique to *B. pilosicoli*. Despite its structural resemblance to bacterial sialidases, BPP43_05035 is a non-catalytic surface lipoprotein that interacts with exposed *N*-glycans on epithelial cells. Interaction and proximity labeling assays in cultured epithelial cells revealed that BPP43-05035 associates directly or closely with proteins that regulate tight junctions (JAM-1, Occludin, and ZO-1) and the actin cytoskeleton (ACTB, MLC2, MYH9, and MYL12B), which fine-tunes tight junction permeability (*21*). Mechanistically, *N*-glycosylation of the extracellular domain of the tight junction protein JAM-1 regulates homophilic dimerization across opposing cells (*25*). Hence, an *N*-glycan-dependent binding of BPP43-05035 to epithelial cells could be mediated via JAM-1, resulting in a weakening of JAM-1 homophilic interactions that subsequently increase tight junction permeability. In line with this argument, BPP43_05035 altered tight junction morphology and increased tight junction permeability to the same extent as viable *B. pilosicoli* in cultured cells, providing critical evidence for the detrimental effect of BPP43_05035-mediated attachment on epithelial monolayers. While changes in tight junction morphology in response to spirochetes have been reported (*26*), we have now identified the bacterial virulence factor that triggers these alterations. In human and murine colon tissue sections, BPP43_05035 bound to the surface glycocalyx and goblet cells, which are compartments rich in *N*-glycosylated mucins. Moreover, our *in vitro* findings were supported by multi-omic analysis of colonic biopsies from non-infected and *B. pilosicoli*-infected participants, where BPP43_05035 was identified in the infected group that displayed transcriptional downregulation of tight junction-associated genes.

Generally, spirochetes are considered part of the human commensal microbiota (*23*), but there is emerging evidence that *B. pilosicoli* and *B. aalborgi* are associated with clinical symptoms including diarrhea and abdominal pain in IBS (*3*), indicating that they are potentially pathogenic bacteria that may trigger symptoms in susceptible individuals. Therefore, it is urgent to develop reliable strategies for the detection of the *Brachyspira* species in the clinic. Due to their strictly anaerobic and slow growth rate, it is difficult to culture *Brachyspira* species (*27*). Instead, the detection of BPP43_05035 in colonic specimens offers opportunities to use the adhesin as a biomarker for the diagnosis and surveillance of *B. pilosicoli* infection. Treatment of *Brachyspira* with antibiotics has raised concerns due to the translocation of bacteria to goblet cells and to the colonic crypts that harbor a vulnerable pool of stem cells (*3*). This dramatic change in host tropism could explain the observed interaction of BPP43_05035 with intracellular proteins that regulate the actin cytoskeleton in epithelial cells. Importantly, our work shows that antibodies against BPP43_05035 can inhibit bacterial attachment to colonic epithelial cells. Consequently, interfering with the intimate attachment of *Brachyspira* to host cells offers an attractive treatment approach to combat infection.

In conclusion, we present BPP43_05035 as a novel adhesin necessary for the attachment of the potentially pathogenic *B. pilosicoli* to human colonic epithelial cells. Based on *in vitro* experiments and analysis of colonic biopsies from human subjects we demonstrate that BPP43_05035 is a species-specific globular protein that deteriorates tight junction integrity *via* engaging host *N*-glycans. Blocking the adhesin-host cell interaction with a specific antibody attenuates the attachment of *B. pilosicoli* to epithelial cells, offering a new therapeutic strategy for the treatment of *B. pilosicoli* infection in humans.

## Acknowledgments

Proteomic analysis was performed at the Proteomics Core Facility, Sahlgrenska Academy, Gothenburg University, with financial support from SciLifeLab and BioMS. We acknowledge the Protein-Glycan Interaction Resource of the Consortium for Functional Glycomics and the National Center for Functional Glycomics (NCFG) at Beth Israel Deaconess Medical Center, Harvard Medical School (supporting grant R24GM137763). We acknowledge the European Synchrotron Radiation Facility (ESRF) for the provision of synchrotron radiation facilities. We also extend our gratitude to the patients and colonoscopists at the Endoscopy Unit of the Sahlgrenska University Hospital.

## Materials and methods

### Human subjects

We recruited participants, healthy controls, and participants with IBS, 18-65 years old through advertisements and referrals to our outpatient clinic specialized in disorders of gut-brain interactions. Participants were screened by questionnaires and clinical interviews to rule out organic gastrointestinal conditions. After providing written informed consent, all participants underwent a sigmoidoscopy without bowel preparation, where biopsies were collected in a fixative (methanol-Carnoy) for future histology and immunohistochemistry, or real-time PCR analysis. Infection with *B. pilosicoli* was confirmed using immunohistochemistry and targeted real-time PCR analysis of biopsy material (*3*). The study was approved by the Gothenburg Ethical Review Board (Ethical number 988-14).

### Mice

C57BL/6N mice of wild-type and Muc2-deficient genotypes were housed in controlled environmental conditions with a temperature range of 21–22°C, a 12-hour light/dark cycle, and *ad libitum* access to food and water. The experiments were approved by the Swedish Laboratory Animal Ethical Committee in Gothenburg, under the ethical permit 2285-19.

### Growth of bacteria

*Brachyspira pilosicoli* strain P43/6/78 (ATCC #51139) was grown in Brain Heart Infusion (BHI) broth (Thermo Fisher Scientific, #CM1135B) containing 10% heat-inactivated fetal bovine serum and 0.2% glucose solution for four days in an anaerobic chamber (Coy Lab Products) at 37°C. *Escherichia coli* strain TUNER cells (Sigma-Aldrich, #70623) were grown in Luria broth (Invitrogen, #12795084) with 30 µg/ml kanamycin at 37°C.

### Mammalian cell culture

Caco-2 (ATCC, #HTB-37) and HCT-116 (ATCC, #CCL-247) cells were cultured in HyClone IMDM Modified medium (Cytiva, #SH30228.01) supplemented with 10% fetal bovine serum (FBS; Thermo Fisher Scientific, #10270106) in a 5% CO_2_ humidified atmosphere at 37°C. Caco-2 cells were cultured for 14-21 days to undergo differentiation and to form a monolayer of polarized cells coupled by tight junctions.

### Recombinant protein production

BPP43_05035, Pal, and tmpB genes were amplified through PCR using specific primers. Subsequently, the amplified DNA was cloned into the pETite N-His vector (Expresso™ T7 cloning and expression system, Lucigen, # 49001-1), resulting in constructs with an N-terminal 6x His tag. Recombinant genes were expressed in the *Escherichia coli* TUNER strain. The bacterial cultures, harboring the respective recombinant plasmids, were cultured until reaching an optical density (OD600) of 0.6 to 0.8. Induction of protein production was achieved by the addition of 0.2 mM isopropyl β-D-1-thiogalactopyranoside (IPTG). Following induction, cells were cultured for an additional 16 hours at 16°C and 180 rpm. Bacterial cells were pelleted, and the resulting pellet was lysed using a lysis buffer composed of 50 mM Tris pH 7.4, 500 mM NaCl, 1X PMSF, 1X cOmplete inhibitor, 10 mM imidazole, and 0.1% IGEPAL. Subsequently, the lysate underwent sonication to clarify the lysate. Purification of the His-tagged proteins was accomplished using Ni Sepharose Excel histidine-tagged protein purification resin (Cytiva, #17371203), and elution was carried out with 350 mM imidazole. An additional purification step was implemented through size exclusion chromatography using a HiLoad 26/600 Superdex 200 pg column (GE Healthcare, # GE28-9893-36) equilibrated with PBS. The concentrations of purified proteins were determined by measuring the absorbance at 280 nm, utilizing the respective molar extinction coefficient for each protein.

### Structure determination and analysis

Crystals of BPP43_05035 were obtained at 18°C by sitting drop vapor diffusion method. The reservoir solution contained 25% polyethylene glycol (PEG) 3350, 0.1 M MgCl_2_, and 0.1 M Bis-Tris buffer at pH 5.2. Single crystals appeared after 5-7 days. Diffraction data were recorded from cryo-cooled crystals (100K) at ESRF beamline ID30B. Data were integrated and merged using XDS (*28*), and scaled, reduced, and further analyzed using CCP4 (*29*) and Coot (*30*). Model refinement was performed with Phenix (*31*). Refinement and data statistics are provided in Table S2. The structure of BPP43_05035 has been deposited with PDB ID 7ZAO. Structural representations were prepared with PyMOL Molecular Graphics System (Schrödinger LLC, Version 4.6) and NGL Viewer (*32*). RMSD calculations were performed in PyMOL using different algorithms depending on sequence similarity. The “Align” command, executing a sequence alignment followed by a structural superposition, was used for proteins with >30% sequence similarity. The “Super” command, which performs a sequence-independent structure-based dynamic programming alignment, was used with proteins with lower sequence identity where “align” algorism failed (NanA, NanI, BPP43_05040, and BPP43_10460). Additional alignments (EPJ79_11040, BHYOB78_13105, and BPP43_01500) were performed using CEalign (*33*) using a combinatorial extension of an alignment path defined by aligned fragment pairs.

### Amino acid sequence analysis

Amino acid sequence alignments were performed using the Clustal Omega multiple sequence alignment tool (*34*).

### Production of a polyclonal antibody against BPP43_05035

Rabbit polyclonal antibody targeting BPP43_05035 was produced by Agrisera antibodies through a multi-step immunization process. Initially, rabbits were immunized once with 200 µg of purified BPP43_05035, without the 6x His tag, mixed with Freund’s complete adjuvant, followed by three subsequent immunizations using 100 µg of BPP43_05035 each time, along with Freund’s incomplete adjuvant. The serum utilized in our investigations was harvested two weeks after the final immunization. As a control, serum collected prior to the first immunization was employed for comparative analyses.

*B. pilosicoli* binding to Caco-2 cells For determining *B. pilosicoli* binding to Caco-2 cells, *B. pilosicoli* were cultured and harvested by centrifugation at 800 x g for 20 min. The pelleted cells were washed three times with sterile, ice-cold PBS. After the final wash, the bacterial cells were suspended in IMDM + 10% FBS and bound on differentiated Caco-2 cells for four hours at 37°C under microaerophilic conditions. The Caco-2 cells were then washed three times with sterile PBS, fixed with 4% paraformaldehyde (PFA) in PBS for ten minutes, permeabilized with 0.1% Triton X-100 in PBS for five minutes, and blocked with 5% FBS in PBS for one hour. Caco-2 cells were washed three times with PBS between the steps and all steps were performed at room temperature (RT). The cells were then incubated with streptavidin Alexa Fluor 488 (Thermo Fisher Scientific, #S11223) and Phalloidin Alexa Fluor 568 (Thermo Fisher Scientific, #A12380) diluted in PBS + 5% FBS overnight at 4°C and washed three times with PBS before mounting and visualizing on a Zeiss LSM700.

Isolation of biotinylated *B. pilosicoli* OMPs For identifying *B. pilosicoli* adhesins involved in Caco-2 cell binding, *B. pilosicoli* cells were pelleted at 800 x g for 20 min and washed as above. Following the final wash, the cell count was adjusted to 25 x 10^6^ cells/ml. A 1 mL suspension of the bacterial cells was mixed with EZ-Link Sulfo-NHS-LC-Biotin (Thermo Fisher Scientific, #A39257) dissolved in 100 µl of sterile MilliQ water. The mixture was incubated at 4°C for 30 minutes. Subsequently, cells were washed three times with ice-cold sterile PBS containing 100 mM glycine. The pellet was resuspended in IMDM + 10% FBS. Bacterial cells were lysed by subjecting them to five 15-second pulses with ten-second intervals using sonication. The lysate was diluted two times and bound onto differentiated Caco-2 cells. The binding reaction was performed for four hours at 37°C. Following binding, Caco-2 cells were washed three times with sterile PBS and then lysed using a lysis buffer (25 mM Tris-HCl, pH 7.4, 150 mM NaCl, 0.1% IGEPAL). The resulting cell lysates were sonicated for ten seconds and centrifuged at 16,000 x g. EZView Red Streptavidin Affinity Gel (Sigma-Aldrich, #E5529) was added to the clarified cell lysate and incubated at 4°C for one hour. The beads were subsequently washed five times with wash buffer (25 mM Tris-HCl, pH 7.4, 150 mM NaCl). To elute bound proteins, 0.1 M glycine at pH 2 was added to the beads. The eluate was neutralized by the addition of 1 M Tris-HCl at pH 8.5. Samples were prepared using filter-aided sample preparation (*35*). Peptides were further cleaned using StageTip C18 columns before subjecting them to mass spectrometry analysis (*36*).

### Binding of bacterial proteins to Caco-2 and HCT116 cells

Fluorescent labeling of BPP43_05035, Pal, and tmpB was achieved using CF488A, CF555, or CF647 dyes, respectively (Biotium, #92213, #92214, #92218), following the manufacturer’s protocol. Fluorescently labeled BPP43_05035, Pal, or tmpB recombinant proteins were diluted in IMDM + 10% FBS and applied to differentiated Caco-2 or HCT116 cells, followed by a two-hour incubation at 4°C. Post-incubation, cells were washed three times with PBS, fixed with 4% PFA in PBS, and blocked with 5% FBS in PBS. The cells were visualized on a Zeiss LSM700.

### Inhibition of BPP43_05035 binding

For blocking endogenous BPP43_05035, *B. pilosicoli* were first labeled with Alexa Fluor 680 (Invitrogen, #A37574) and then subjected to a one-hour pre-incubation with a 1:25 dilution of anti-BPP43_05035 or control serum at RT. Following pre-incubation, *B. pilosicoli* were added to Caco-2 cells for four hours at 37°C under microaerophilic conditions. For competitive inhibition of BPP43_05035 binding sites on differentiated cells, Caco-2 cells on a 96-well plate were pre-incubated with 10 µM BPP43_05035 for two hours at 4°C. After the pre-incubation, Alexa Fluor 680-labeled *B. pilosicoli* were added to Caco-2 cells for four hours at 37°C under microaerophilic conditions. Subsequently, cells were washed with PBS, fixed with 4% PFA, and analyzed using an Odyssey CLx near-infrared fluorescence imaging system (LI-COR Biosciences) or a Zeiss LSM700 microscope. All image analysis and processing were performed using Image Studio (LI-COR Biosciences) and ImageJ software v.1.53t (National Institutes of Health, Bethesda, MD).

### Antibody staining of *B. pilosicoli*

*B. pilosicoli* cultures were washed twice with PBS. The resulting washed cell suspension was fixed with 4% PFA for ten minutes at RT, followed by a single wash with PBS. Subsequently, bacterial cells were incubated overnight at 4°C with either a 1:5000 dilution of anti-BPP43_05035 or control serum. After three washes with PBS, the cells were stained using anti-rabbit IgG Alexa Fluor 488 (Invitrogen, #A21206) at a 1:300 dilution for two hours at RT. Following additional washes, the cells were mounted on slides using Prolong Gold antifade (Thermo Fischer Scientific, #P36980). Imaging was performed on a Zeiss LSM900, equipped with an AiryScan2 detector and a plan-Apochromat 63x/1.4 Oil DIC M27 lens.

### Immunohistochemistry

Sigmoid colon biopsies obtained from participants or mouse distal colon were fixed with Carnoy’s fixative, comprising 60% absolute methanol, 30% chloroform, and 10% glacial acetic acid. The fixed tissues were then embedded in paraffin, and paraffin-embedded sections were subjected to deparaffinization with a xylene substitute at 60°C, followed by rehydration through a series of ethanol concentrations (100%, 70%, 50%, and 30%). Tissue demarcation was established using the ImmEdge Hydrophobic Barrier PAP pen (Vector Laboratories, #H-4000), followed by permeabilization with 0.1% Triton X-100 and blocking with 5% FBS in PBS. Subsequently, the samples were incubated overnight at 4°C with 5 µM BPP43_05035 diluted in 5% FBS in PBS. After three washes, sections were exposed to anti-BPP43_05035 (1:5000) and anti-ezrin antibody (1:500; Sigma-Aldrich, #E8897), overnight at 4°C. Following three PBS washes, sections were treated with Alexa Fluor 488 goat anti-rabbit (1:1000; Invitrogen, #A11055) and Alexa Fluor 555 donkey anti-mouse (1:1000; Invitrogen, #A31570). Nuclear staining with Hoechst 33258 (1:10,000; Sigma-Aldrich, #94403) was performed for five minutes at RT, and coverslips were mounted for imaging using a Zeiss LSM 700.

### Immunocytochemistry

Caco-2 cells were cultured and stained on chamber slides (Thermo Fisher Scientific, #154534PK). Cells were washed 3 times in PBS followed by a 10-minute fixation in 4% PFA in PBS at RT. Excess PFA was washed away with three PBS washes and cells were permeabilized by 0.1% Triton X-100 in PBS for five minutes. After permeabilization, cells were washed three times with PBS and blocked for one hour in 5% FBS in PBS at RT. Subsequently, the samples were incubated overnight at 4°C with 5 µM BPP43_05035 diluted in 5% FBS in PBS. After three washes, cells were exposed to anti-BPP43_05035 (1:5000) and anti-ezrin antibody (1:500) overnight at 4°C. Tight junctions were stained after BPP43_05035 binding with 1:500 dilutions of anti-JAM-A (1:500; Sigma-Aldrich, #SAB4200468), anti-Occludin (1:500; Thermo Fischer Scientific, #33-1500) and anti-ZO-1 (1:500; Thermo Fischer Scientific, #33-9100).

Following three PBS washes, cells were treated with Alexa Fluor 488 goat anti-rabbit (1:1000) and Alexa Fluor 555 donkey anti-mouse (1:1000). Nuclear staining with Hoechst 33258 (1:10,000) was performed for five minutes at RT, and coverslips were mounted for imaging using a Zeiss LSM 700. To cleave *N*-glycans from the cells, *N*-Glycosidase F (Roche, #11365185001) was diluted 1:10 in serum-free IMDM and incubated with the cells for two hours at 37°C prior to fixation, permeabilization, and blocking.

### SDS-PAGE, immunoblots, and protein gels

Proteins were subjected to reduction in 4X reducing sample buffer (8% SDS, 400 mM Dithiothreitol) and resolved on a precast 4%–12% SDS-polyacrylamide gel (Thermo Fisher Scientific, #XP04125BOX). Coomassie staining was performed using Imperial Protein Stain (Thermo Fisher Scientific, #24615) for one hour at RT, followed by destaining in water, and visualization on an Odyssey CLx near-infrared fluorescence imaging system (LI-COR Biosciences). Protein transfer to a PVDF-FL membrane (Millipore, #05317) was achieved at a current of 2.5 mA/cm^2^ for one hour. The membrane was then blocked in 5% FBS in PBS for one hour and incubated overnight at 4°C with *Brachyspira* antiserum (1:5000) (*11*) in 5% FBS in PBS + 0.1% Tween-20 (PBS-T). After three washes in PBS-T, the membrane was incubated with goat anti-rabbit Alexa Fluor 790 (1:20,000; Invitrogen, #A11369) secondary antibody or streptavidin Alexa Fluor 680 (1:20,000; Invitrogen, #S21378) for one hour at RT in the dark. After three washes in PBS-T, the membrane was visualized on an Odyssey CLx near-infrared fluorescence imaging system (LI-COR Biosciences).

### Thin layer chromatography

For the evaluation of sialidase activity of BPP43_05035, 10 μM of BPP43_05035 was subjected to incubation with 0.5% porcine gastric mucin II or III (Merck, #M2378 and #M1778) in reactions conducted under pH 5.5-8.5, over 16 hours at 37°C. Subsequently, 2 μL of each sample was applied to thin layer chromatography silica plates and separated using a butanol:acetic acid:water (2:1:1) running buffer. The resulting TLC plates were air-dried, and visualization of released sugars was achieved through diphenylamine staining, involving a solution composed of 1 ml of 37.5% HCl, 2 ml of aniline, 10 ml of 85% H_3_PO_3_, 100 ml of ethyl acetate, and 2 g of diphenylamine, followed by heating at 100°C for 20 minutes.

### Hydrolysis assays

For the evaluation of esterase activity of BPP43_05035, 1 µM or 10 µM of BPP43_05035 was incubated with 1 mM of 4-nitrophenyl acetate, 4-Nitrophenyl butyrate, and 4-Nitrophenyl octanoate (Sigma-Aldrich, #N8130, #N9876, #21742) (pH 5.5-8.5) at 30°C for one hour. Production of p-nitrophenol was measured at 410 nm.

### Glycan array

The glycan array analysis was conducted using 5 µg/ml, 50 µg/ml, or 500 µg/ml concentrations of Alexa Fluor 488 labeled-BPP43_05035 against a repertoire of 562 unique glycan structures, spotted on a CFG Glycan Array version 5.5. This experimental procedure was executed by the National Center for Functional Glycomics according to a standard protocol ” Direct Glycan Binding Assay for Fluorescent Labeled Sample”, accessible at https://research.bidmc.org/ncfg/protocols. Glycan structures were drawn according to Symbol Nomenclature For Glycans (SNFG) guidelines, using DrawGlycan-SNFG (*37*).

### Trans-epithelial electrical resistance (TEER) measurement

Caco-2 cells underwent differentiation on Transwell filters (Merck, #CLS3496). Before the experiment, cells were washed with IMDM +10% FBS, followed by incubation with fresh media for one hour at 37°C. Transepithelial electrical resistance (TEER) was measured using a manual Epithelial Volt Ohm Meter (EVOM2; World Precision Instruments), after which 100 µM BPP43_05035, diluted in IMDM +5% FBS, was applied to the cells. This was followed by a four-hour incubation at 37°C in a 5% CO_2_ humidified atmosphere. After the four-hour incubation, TEER was measured again. For TEER assessments involving viable *B. pilosicoli*, fresh media was added, and cells were incubated for one hour at 37°C under microaerophilic conditions. Subsequently, *B. pilosicoli* was introduced, and TEER was immediately measured. Additional TEER readings were obtained after four hours, 24 hours, and 30 hours under microaerophilic conditions.

### Proximity labeling by antibody recognition

3 x 10^5^ Caco-2 cells were seeded on 9.6 mm^2^ wells and differentiated for 14 days. *B. pilosicoli* were grown to mid-log phase (15-16 hrs) and harvested by centrifugation at 800 x g for 20 min at RT. Bacterial cells were gently resuspended in cell culture medium and the OD was adjusted to 0.9 corresponding to 1 x 10^10^ bacterial cells/mL. Caco-2 monolayers were seeded with 7 x 10^9^ bacterial cells for 2 hours at 37°C under microaerophilic conditions. Cell monolayers were gently washed three times with PBS, fixed with 4% paraformaldehyde in PBS for 10 min at RT, and washed twice in PBS-T. Cells were permeabilized in PBS + 0.5% Triton X-100 for 10 min at RT followed by 3 x 10 min washes with PBS-T. 30 mM H_2_O_2_ in PBS was added overnight at RT to quench the endogenous peroxidase activity. 30 mM fresh H_2_O_2_ in PBS was added for 30 min followed by two 10 min washes with PBS-T. Cells were incubated with a blocking buffer (5% BSA in PBS) for 2 hours on an orbital shaker. Cells were stained with *Brachyspira* antiserum (1:2000) diluted in blocking buffer overnight at 4°C in a humid chamber on an orbital shaker. After three subsequent 1-hour PBS-T washes, cells were incubated with HRP-conjugated goat anti-rabbit secondary antibody (1:1000; Sigma-Aldrich, # 12-348) diluted in blocking buffer for 1 hour at RT. Unbound antibody was removed by 3 x 2-hour washes with PBS-T and the cells were pre-incubated with a final concentration of 500 μM biotin-XX-tyramide (Sigma-Aldrich, # SML3484) at RT. After 10 min, a final concentration of 2.5 mM H_2_O_2_ in PBS was added to the biotin tyramide solution for 1 min at RT. To quench the reaction, 500 μL of 500 mM sodium ascorbate (3 x 5 min) was added to the cells followed by 3 x 10 min washes with PBS-T. Cells were lysed in 200 μL PBS-T + 2% SDS + 2% deoxycholate + 1X complete protease inhibitor. Cells were collected by scraping, sonicated for 10 seconds, and boiled at 95°C for 60 minutes. Samples were cleared by centrifugation at maximum speed for 10 min. The supernatants were diluted in 1 mL PBS-T and 50 μL was saved as an input control. 20 μL of pre-washed Streptavidin Dynabeads (Thermo Fisher Scientific, #11205D) was added to each sample and incubated for 48 hours, rotating at 4°C. Beads were washed in 1 x 15 mL PBS-T, 1 x 15 mL PBS-T + 1M NaCl, 1 x 15 mL PBS, and 1 x 15 mL PBS + 0.5% Triton X-100. Beads were re-suspended in PBS and proteins were eluted by adding 1 volume of 2X lysis buffer (4% SDS, 200 mM DTT, 125 mM Tris HCl pH 6.8) and boiling at 95°C for 5 min. The eluate was separated from beads using a DynaMag-2 magnet (Thermo Fisher Scientific, #12321D).

### Co-immunoprecipitation

3 x 10^5^ Caco-2 cells were seeded on 9.6 mm^2^ wells and differentiated for 14 days. Cells were lysed in lysis buffer (25 mM Tris-HCl, 150 mM NaCl, 1% Triton X-100, pH 7.4) containing 1X protease inhibitor cocktail for 20 min on ice. Cells were scraped into a 1.5 mL Eppendorf tube and sonicated 3 x 30 seconds on 40% amplitude in a water-submerged sonicator at 4°C. Lysates were cleared by centrifugation at 16,000 x g for 30 min at 4°C. 100 µg of purified recombinant BPP43_05035 was incubated with 1 mL of cell lysate overnight at 4°C. 10 µL of pre-serum or BPP43_05035 specific antibody was added to each tube, which was then incubated overnight at 4°C. 50 µL of Dynabeads™ M-280 Sheep Anti-Rabbit IgG (Thermo Fisher Scientific, #11203D) was added to each tube followed by incubation at 4°C for 2 hours. Dynabeads were isolated using a DynaMag-2 magnet and washed in 3 x 1 mL Lysis buffer for 5 min at 4°C. Bound proteins were eluted using a reducing sample buffer.

### RNA sequencing

RNA was extracted from whole biopsies from the sigmoid colon using an RNeasy Mini kit (Qiagen, #74106). The quality of the isolated RNA was determined using a Bioanalyzer (Agilent) with a RIN value greater than 8. cDNA libraries were prepared using the TrueSeq Stranded Total RNA Sample Preparation kit with Ribo-Zero Plus (Illumina) according to the manufacturer’s protocol and sequenced thereafter via paired-end with the NovaSeq 6000 platform (Illumina). The sequencing quality was measured for all lanes, reads, and cycles with 93.45% of bases above Q30. The quality of raw reads was assessed using FastQC (version 0.11.2) evaluating for per base sequence quality and adaptor contamination. The reads were mapped against the human transcriptome, release 38 (GRCh38.p13) from Gencode using Salmon (version 1.10.1) (*38*). The expression levels of each gene were calculated as transcripts per million (TPM) using R library tximport (v1.18.0) (*39*). The read counts were quantified and normalized by the median of ratio method via the R package DESeq2 (v1.30.1) (*40*). Heat map and PCA plot of human transcriptome were generated using ClustVis (*41*).

### Proteomic analysis after proximity labeling and co-immunoprecipitation

Samples from proximity labeling were centrifuged at 16,000 x g at RT and the supernatant was added onto 10 kDa cutoff filters (PALL Cat# OD010C33). Proteins were digested using filter-aided sample preparation with sequence-grade trypsin (Promega, # V5111) at 37°C overnight. Peptide concentration after elution was measured at 280 nm using NanoDrop (Thermo Fisher Scientific) and peptides were cleaned with StageTip C18 columns (67) before mass spectrometry (MS). For proteomic analysis after co-immunoprecipitation, protein gels were stained with Imperial Protein Stain. The bands to be analyzed were cut out from the gel, the gel pieces were shaken three times for 40 min. in 25 mM NH_4_HCO_3_ and 50% acetonitrile and dried by vacuum centrifugation. Samples were incubated with sequence-grade trypsin (10 μg/mL in 25 mM NH_4_HCO_3_) for 16 hours at 37°C. Peptides were eluted from the gel pieces by shaking in 5% TFA with 75% acetonitrile for 30 min., the eluate collected, and the gel pieces were washed again 5% TFA with 75% acetonitrile for 30 min.. The two fractions were pooled, dried by vacuum centrifugation, and dissolved in formic acid (0.1%). Peptides were cleaned with StageTip C18 columns before mass spectrometry.

Nano LC-MS/MS was performed on a Q-Exactive HF mass-spectrometer (Thermo Fischer Scientific), connected with an EASY-nLC 1000 system (Thermo Fischer Scientific) through a nanoelectrospray ion source. Peptides were loaded on a reverse-phase column (150 mm x 0.075 mm inner diameter, New Objective, New Objective, Woburn, MA), packed in-house with Reprosil-Pur C18-AQ 3 mm particles (Dr. Maisch, Ammerbuch, Germany). Peptides were separated with a 230-minute gradient: from 3% to 25% B in 175 min., 25% to 45% B in 30 min., 45% to 100% B in 5 min., followed by 20 min. wash with 100% of B (A: 0.1% formic acid, B: 0.1% formic acid/80% acetonitrile) using a flow rate of 250 nl/min.. Q-Exactive HF was operated at 250°C capillary temperature and 2.0 kV spray voltage. Full mass spectra were acquired in the Orbitrap mass analyzer over a mass range from m/z 350 to 1600 with a resolution of 60,000 (m/z 200) after the accumulation of ions to a 3e6 target value based on predictive AGC from the previous full scan. For each full MS scan, the twelve most intense peaks with a charge state ≥ 2 were fragmented in the HCD collision cell with a normalized collision energy of 27%. The tandem mass spectrum was acquired in the Orbitrap mass analyzer with a resolution of 15,000 after the accumulation of ions to a 1e5 target value. Dynamic exclusion was set to 30 seconds. The maximum allowed ion accumulation times were 20 ms for full MS scans and 50 ms for tandem mass spectrum. MS raw files were analyzed with MaxQuant software version 1.5.7.4. Peak lists were identified by searching against the reviewed database of Homo sapiens (Uniprot ID: UP000005640, 20361 entries). Searches were performed using trypsin as an enzyme, maximum of 2 missed cleavages, and a precursor tolerance of 20 ppm in the first search used for recalibration, followed by 7 ppm for the main search and 0.5 Da for fragment ions. Carbamidomethylation of cysteine was set as a fixed modification, and methionine oxidation and protein N-terminal acetylation were set as variable modifications. The required false discovery rate (FDR) was set to 1% for peptide and protein levels, and the minimum required peptide length was set to seven amino acids.

### Proteomic analysis of human biopsies

Frozen biopsies were thawed on ice, and PBS was added to the tissue, followed by rotation of the sample for five minutes at 4°C. Subsequently, the PBS was aspirated, and cell recovery solution (Corning, #CLS354253) was added, with the sample rotated at 4°C for 15 minutes. After a 30-second vortex, the remaining tissue was removed from the solution, and the sample was centrifuged at 800 rpm for five minutes at 4°C. Cell pellets were lysed in 2% sodium dodecyl sulfate and 50 mM triethylammonium bicarbonate (TEAB) using a Covaris ML230 ultrasonicator. Protein concentrations were determined using Pierce BCA Protein Assay Kit (Thermo Scientific) on a Benchmark Plus microplate reader (BIO-RAD). 15 µg protein amount of every sample was reduced in 10 mM dithiothreitol at 56°C for 30 min and alkylated in 20 mM iodoacetamide at room temperature for 30 min. The samples were mixed with hydrophobic and hydrophilic Carboxylate-Modified Sera-Mag™ SpeedBeads (Cytiva, #GE44152105050250 and 45152105050250) with a bead-to-protein ratio of 10:1. The SP3 workflow was adapted from the protein and peptide clean-up for mass spectrometry protocol provided by the manufacturer. In short, proteins were precipitated on the beads with 100% ethanol, washed with 80% ethanol, and dried at room temperature. Beads were resuspended in 50 mM TEAB, and proteins were digested with LysC and trypsin (Promega) in two steps with a final enzyme-to-protein ratio of 1:25. Beads were removed, and peptides were labeled using TMTpro 18-plex isobaric mass tagging reagents (Thermo Fisher Scientific, #A52045) according to the manufactureŕs instructions. The labeled samples were pooled into one TMT set. The TMT set was purified using a HiPPR detergent removal kit (Thermo Fisher Scientific, # 87780) and Pierce peptide desalting spin columns (Thermo Fisher Scientific, #89851), according to the manufactureŕs instructions. The TMT-set was fractionated by basic reversed-phase chromatography using a Dionex Ultimate 3000 UPLC system (Thermo Fischer Scientific) equipped with a reversed-phase XBridge BEH C18 column (3.5 μm, 2.1×150 mm, Waters Corporation). Peptides were eluted with a stepped gradient from 3% to 80% solvent B over 65 min followed by an increase to 80% B at a flow of 200 µL/min. Solvent A was 25 mM ammonia and solvent B was 84% acetonitrile. 96 primary fractions were combined into 30 final fractions which were evaporated and reconstituted in 3% acetonitrile, and 0.1% trifluoroacetic for LC-MS3 analysis.

The fractions were analyzed on an Orbitrap Fusion Lumos Tribrid mass spectrometer equipped with a FAIMS Pro ion mobility system and interfaced with an Easy-nLC1200 liquid chromatography system (all Thermo Fisher Scientific). Peptides were trapped on an Acclaim Pepmap 100 C18 trap column (100 μm x 2 cm, particle size 5 μm, Thermo Fisher Scientific) and separated on an in-house packed analytical column (35 cm x 75 μm, particle size 3 μm, Reprosil-Pur C18, Dr. Maisch) using a stepped gradient from 5% to 35% acetonitrile in 0.2% formic acid over 77 min at a flow of 300 nL/min. FAIMS Pro was alternating between the compensation voltages (CV) of -50 and -70, and the same data-dependent settings were used at both CVs. The precursor ion mass spectra were acquired at a resolution of 120,000 (m/z 200) and an m/z range of 375-1500. Using a cycle time of 1.5 seconds the most abundant precursors with charges 2-7 were isolated with an m/z window of 0.7 and fragmented by collision-induced dissociation (CID) at 30%. Fragment spectra were recorded in the ion trap at a Rapid scan rate. The ten most abundant MS2 fragment ions were isolated using multi-notch isolation for further MS3 fractionation. MS3 fractionation was performed using higher-energy collision dissociation (HCD) at 55% and the MS3 spectra were recorded in the Orbitrap at 50,000 resolution and an m/z range of 100–500.

Data analysis was performed using Proteome Discoverer version 3.0 (Thermo Fisher Scientific) and Sequest HT as a search engine. The data was matched against the reviewed database of Homo sapiens (Uniprot ID: UP000005640, 20361 entries) and *B. pilosicoli* P43/6/78 (Uniprot ID: UP000010793, 2208 entries) allowing 2 missed cleavages. Precursor mass tolerance was set to 10 ppm and fragment mass tolerance was set to 0.6 Da. Cysteine carbamidomethylation and TMTpro were set as fixed modifications, while methionine oxidation and protein N-terminal acetylation were set as variable modifications. Percolator was used for PSM validation at an FDR of 1%. The abundance values were normalized to the total peptide amount.

### KEGG pathways and Gene ontology analysis

KEGG pathway and gene ontology (GO) analyses were performed using the String database (*42*). The full STRING network was queried for the identified proteins with a medium confidence of 0.400. Analysis output was KEGG pathway and GO: Cellular component.

### Quantification and statistical analysis

Data analysis was performed using GraphPad Prism (version 9.5). The unpaired t-test with Welch’s correction, assuming non-equal SDs was used for the comparison of two groups. *p< 0.05, **p<0.01, ***p<0.001, ****p<0.0001.

