## Supplemental material for "BPP43_05035 is a *Brachyspira pilosicoli* cell surface adhesin that weakens the integrity of the epithelial barrier during infection"

**Figure S1. Sequence and structural homologs of BPP43_05035 identified by DALI.**

**(A)** Ten top hits generated by BLAST search of the BPP43_05035 amino acid sequence.

**(B)** Eight top hits generated by DALI search of the BPP43_05035 crystal structure.

**(C)** Structural alignment and projections of BPP43_05035 (7ZAO) (brown), NanA (2AY7) (blue), and NanI (2BF6) (green).

**Figure S2. Hydrolysis assay using 4-nitrophenyl esters as substrates at pH 7.**

**(A)** A cartoon representation of the putative catalytic triad in BPP43_05035. Histidine (H) 221, glutamate (E) 282, and serine (S) 340 are depicted.

**(B)** Activity of BPP43_05035 (10 μM) against 0.5% of PGM type II and III across pH 5.5-8.5. Arrowheads point to sialic acid used as a standard (Std).

**(C)** A cartoon representation showing the overlay of *Vibrio cholerae* sialidase NanH (PDB ID 1W0P) (blue) and BPP43_05035 (PDB ID 7ZAO) (brown). Residues in the catalytic site of NanH are overlayed with the residues in the putative catalytic site of BPP43_05035.

**(D)** Activity of BPP43_05035 (0-10 μM) against 1 mM 4-nitrophenyl acetate across pH 5.5-8.5. n=2 for each group. Data are means ± SD. Significance at each pH was determined by unpaired t-test.

**(E)** Activity of BPP43_05035 (10 μM) against (10 nM-1 µM) 4-nitrophenol, 4-nitrophenyl butyrate, and 4-nitrophenyl octanoate at pH 7.0.
